# A pragmatic reevaluation of the efficacy of nonhuman primate optogenetics for psychiatry

**DOI:** 10.1101/2020.12.10.420331

**Authors:** Eliza Bliss-Moreau, Vincent D. Costa, Mark G. Baxter

**Affiliations:** Department of Psychology, University of California Davis; California National Primate Research Center, University of California Davis; Department of Behavioral Neuroscience, Oregon Health Sciences University; Oregon National Primate Research Center, Oregon Health Sciences University; Nash Family Department of Neuroscience, Icahn School of Medicine at Mount Sinai

**Keywords:** optogenetics, nonhuman primates, translational neuroscience, translational psychiatry, open science

## Abstract

Translational neuroscience is committed to generating discoveries in the laboratory that ultimately can improve human lives. Optogenetics has received considerable attention because of its demonstrated promise in rodent brains to manipulate cells and circuits. In a recent report, Tremblay and colleagues (2020) introduce an open resource detailing optogenetic studies of the nonhuman primate (NHP) brain and make robust claims about the translatability of the technology. We propose that their quantitative (e.g., a 91% success rate) and theoretical claims are questionable because the data were analyzed at a level relevant to the rodent but not NHP brain, injections were clustered within a few monkeys in a few studies in a few brain regions, and their definitions of success was not clearly relevant to human neuropsychiatric disease. A reanalysis of the data with a modified definition of success that included a behavioral *and* biological effect revealed an 62.5% success rate that was lower when considering only strong outcomes (53.1%). This calls into question the current efficacy of optogenetic techniques in the NHP brain and suggests that we are a long way from being able to leverage them in “the service of patients with neurological or psychiatric conditions” as the Tremblay report claims.

---

The goal of translational neuroscience is to generate discoveries at the bench that can be implemented at the bedside, ultimately improving people’s lives. Yet traversing the landscape between bench and bedside to forge a translational bridge is so challenging that it is often referred to as the valley of death (Insel, 2011). Robust, repeatable, valid methods and experiments carried out in cell lines and some animal models often fail to produce tractable observations, treatments, and interventions for humans. These failures occur for many reasons but one of the most prominent and important reasons is that the most frequently used models, be they cell lines or animal models, do not recapitulate the human brain’s structure and function with high enough fidelity to allow for translation. Dominant animal models, like mice, do not share key neuroanatomical, developmental, or behavioral homologies with humans (Laubach et al., 2018; Preuss & Wise, 2022; Wallis et al., 2017; Wise, 2008). Yet many nonhuman primates, particularly cercopithecine monkeys such as macaques, do share such homologies and thus can be a translational bridge between bench and bedside (e.g., Roberts & Clarke, 2019; Rudebeck et al., 2019).

Nowhere is there more demand to traverse the *translational neuroscience valley of death* than in the case of new technology that allows for precise modulation of cellular level activity within the brain. Tools like chemogenetics, in which a viral vector is used to transfect neurons so that they express a designer receptor which responds to a specific chemical compound, and optogenetics, in which a viral vector is used to implant light-sensitive ion channels in cell membrane that then can be activated/deactivated with a light, have been suggested to have the capacity to radically change the health landscape for humans by generating new treatments and interventions to modulate cells, circuits, and ultimately human behavior and experience. Evaluating the efficacy of these newer technologies is critical because the tradeoff between technologies is a zero-sum game in the research funding landscape – once new tools are thought to be effective, grant applications using better-established technologies tend to be criticized as lacking innovation. Moreover, the rush to use new technology for technology’s sake may deplete scientific resources when the failure rate of new techniques is high.

Originally developed in mice, optogenetics is being increasingly used in nonhuman primates over the last few years, promising the use of circuit-specific modulation via light in the primate brain (El-Shamayleh & Horwitz, 2019). The goal of this commentary is to explore the idea that nonhuman primate optogenetics is a functioning technology for basic neuroscience, let alone for translational neuroscience with clinical applications as has been recently claimed. We focus on a recent review of published and unpublished optogenetics studies in nonhuman primates, recently compiled into an open resource (Tremblay et al., 2020).

## The evolution of technology and the hype cycle

All technological development occurs in cycles, and the development of neuroscientific tools is no different. One useful model for understanding technology development is “the hype cycle” (see Figure 1A) which details the evolution of a technology over time from the point of idea conception to the point at which the technology becomes truly productive (Blosch & Fenn, 2018). Five phases emerge when time (x-axis) is plotted relative to “expectations” for the new technology (y-axis). During the initial “innovation trigger” a breakthrough of some sort sparks interest in the new technology. The innovation trigger is followed by an increase in expectations that leads to the “peak of inflated experiences” – this building wave of expectations occurs because of, and also represents, the excitement about the new technology and its promise. At some point, expectations are not met because they exceed the current capacity of the technology – this is the peak. Hype about the technology spreads and people attempt to outcompete each other to use it, despite the fact that the technology is likely to be immature and its efficacy is either unclear or demonstrably low. Once the community realizes that the expectations exceed capacity and that there are major hurdles to deploying the technology, there is a dramatic decrease in expectations down to the “trough of disillusionment”. From here, a dedicated few carry out explicit work to address the hurdles and the mismatch between expectations and capacity and the technology matures, ultimately leading to the “slope of enlightenment” in which is characterized by a developing understanding about the contexts in which the technology is valuable and the contexts in which the technology is not valuable. Finally, the cycle ends with the “plateau of productivity” which follows an increase of technology adoption as real-world benefits are realized, although the level of the plateau (and therefore the expectations realized by the technology) varies across technologies. Throughout this entire cycle, the potential users of the technology are trying to optimize their use of it relative to its maturity and efficacy so that they do not adopt the technology too early (and thus waste resources) or too late to be competitive, nor do they give up on the technology too soon or continue using it for too long.

**Figure 1.**
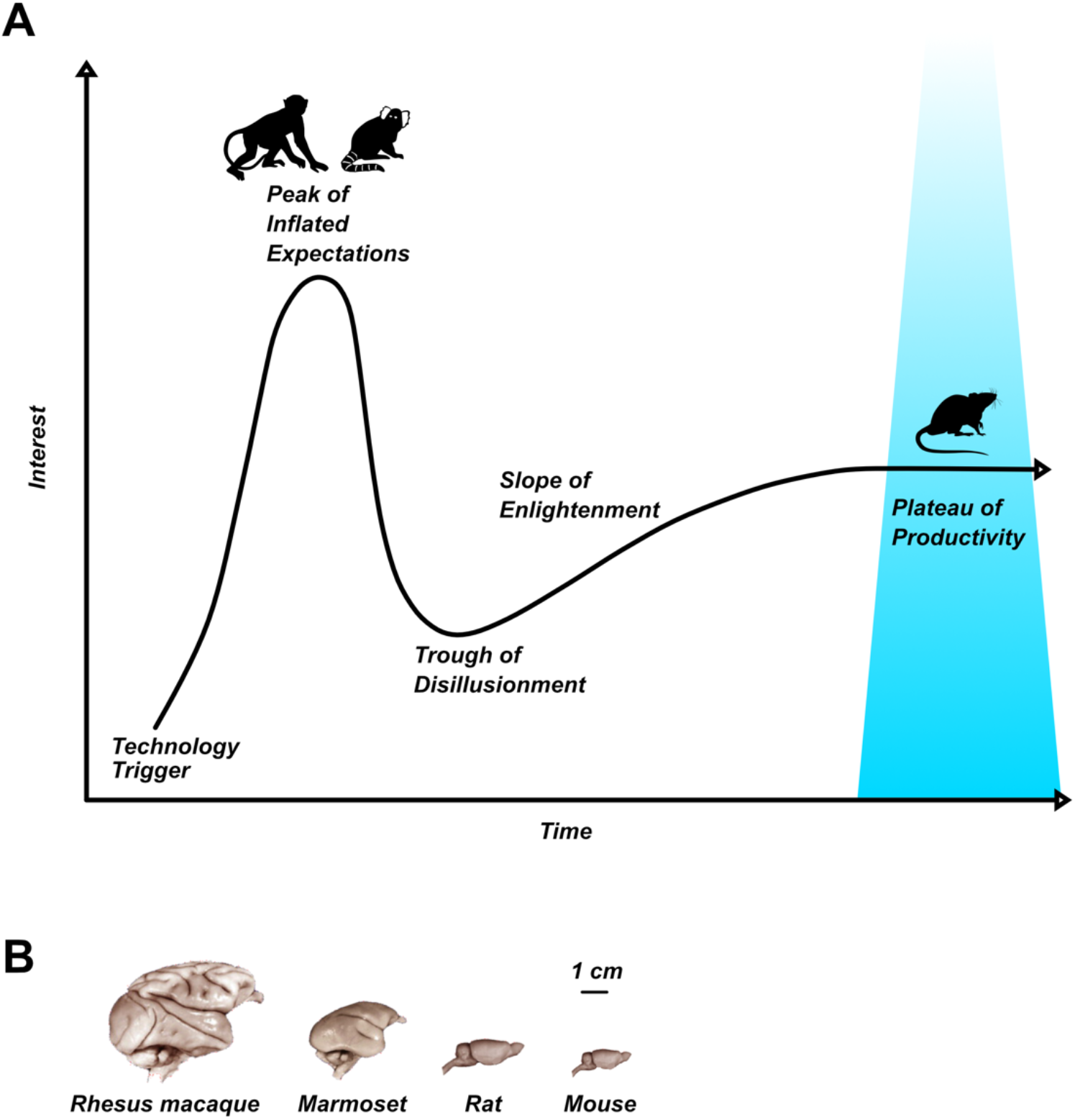
Optogenetics as a technology is in different phases of the hype cycle depending on the species in which it is deployed (A) in part because the brains of research species differ in terms of size and complexity (B) with nonhuman primate brains (rhesus macaque, marmoset) being significantly bigger and more complex than rodent brains (rat, mouse).

From a translational science perspective, a neurobiological technology is only truly *useful* if it can be readily deployed in humans and meets expectations in terms of addressing health concerns when it is deployed – the technology is in the plateau of productivity. What this would mean for optogenetics is that human neurons could be transduced, a light could be placed near the neurons (ideally without causing significant damage to other brain areas), and then activation or deactivation of those neurons via light would alter neural activity and behavior or experience.

Optogenetics, like many neurobiological technologies, was developed in mice who have radically different brains than humans. Optogenetics appears to be in the plateau of productivity in murine brains (Deisseroth, 2015; Fenno et al., 2011), especially in combination with genetic engineering to facilitate targeted, cell-type specific expression of opsins. But differences in mouse and human behavior and biology call into question the efficacy of this technology for human application. First, lissencephalic mouse brains are structured such that many of the important neural hubs for behavior, generally, and for neuropsychiatric disease, specifically, are on the surface of the brain which means that they are easier to access both with the virus to transfect neurons and the light needed to modify neural activity (Figure 1B). Second, while some behavioral assays in mice may exhibit face validity with specific features of behaviors associated with human neuropsychiatric diseases, the extent to which they capture the complexity of the behaviors associated with human neuropsychiatric diseases and other forms of validity (e.g., construct, predictive) is questionable. Aberrations in complex behaviors leading to depression, anxiety, compulsions, and psychosis are likely emergent and rely on degenerate circuitry (for a discussion of degeneracy see Edelman & Gally, 2001). Thus, explicit validation of the effectiveness of optogenetic manipulation of brain circuits relevant to psychiatry in larger NHP brains is doubly critical, in terms of the physical properties of the brain as well as the modulation of more complex behavioral processes.

## Has nonhuman primate optogenetics reached the plateau of productivity?

The goal of translating optogenetics to humans has generated increased interest in carrying out optogenetic studies in nonhuman primates who are thought to be a translational bridge for the valley of death. Enough studies using optogenetics in nonhuman primates have been carried out and/or published to now allow for the concatenation of an open resource to share nonhuman primate optogenetics data (https://osf.io/mknfu/). A companion manuscript was published by Tremblay et al (2020) detailing those data and making, in our opinion, bold claims about what the implications of their collected data for the readiness of optogenetics to remediate neuropsychiatric illness. For example, these authors state “we hope this resource will be used not only by basic scientists trying to uncover the workings of the primate brain using optogenetics but also by translational scientists hoping to bring this powerful technology to the service of patients with neurological or psychiatric conditions” (p 12). This suggests that the open resource has direct translatability and the technology is ready to be deployed for the treatment and intervention of neuropsychiatric diseases. Such uses would suggest that primate optogenetics is in the “plateau of productivity” of the hype cycle, or at the very least further along the x-axis than the “trough of disillusionment”.

The open resource (Tremblay et al., 2020) reflects contributions from 45 research laboratories, with 1042 individual data points included (552 previously unpublished) representing experiments in 198 monkeys and thus aggregates an enormous amount of technical expertise on neurosurgical placement of viral vectors in the NHP brain, a substantial contribution to the field. Moreover, meta- and open science efforts in NHP neuroscience are critical and until very recently, very rare. By metascience, we refer to a collection of approaches that include meta-analyses (analyses of existing analyses), pooling of data across laboratories, splitting data collection efforts strategically across laboratories to guard against protocols being laboratory specific, and gathering published and unpublished data in shared resources, among others (e.g., the “many labs” projects; e.g., (Ebersole et al., 2016; Klein et al., 2014, 2018). These efforts are important for NHP neuroscience because our field typically uses small sample sizes and has had a historical bias against replicating experiments, constraints that are both ethical and practical. By open science, we mean the sharing of scientific resources including primary materials and data (for an NHP specific example see (Milham et al., 2020; Milham et al., 2018). Combining meta- and open data practices, as the optogenetics database does, sets the stage for scientists to be able to evaluate the effectiveness of methods or experiments and to determine where efforts are needed to move science forward and to determine the rigor and reproducibility of the established methods (see Bliss-Moreau et al., 2021 for a discussion). This particular effort deserves significant recognition because of the large number of unpublished data sets and the number of “unsuccessful” studies included in it. These two types of data are critical for determining if an approach is truly effective. Without inclusion of those types of data, particularly in the context of a publication culture that focuses heavily on null hypothesis testing and rewards significant effects to a greater degree than those that are not significant (with regards to ease and visibility of publications), the success of an approach is likely to be overestimated. In that vein, efforts that gather data that is both published and unpublished, as well as successful and unsuccessful, on the efficacy of new tech is both important and laudable.

The large database of optogenetics studies in NHPs positions us well to evaluate where optogenetic technology in NHPs is along the hype cycle and whether or not it is ready for deployment in a translational context – either to ask questions about the etiology of neuropsychiatric diseases or to actually be deployed to treat such diseases. In reading Tremblay et al. (2020), we drew different conclusions about the usefulness of optogenetics in NHPs and in turn the impact of this method in the service of understanding and developing treatment interventions for psychiatric and neurologic disease. Two issues are notable. First, the definition of experimental success in Tremblay et al. (2020) was overly broad on two dimensions. Studies were considered successful if they generated histological, *or* neurophysiological, *or* behavioral effects, and what was counted as a successful behavioral effect included subjective evaluations of narrative accounts of animals’ behaviors as well as effects qualitatively judged to be “weak”. At a minimum, the successful use of optogenetics in behavioral or systems neuroscience (separate from its translational relevance) would require some evidence that neurons have been modulated either via histology of viral expression or altered neuronal activity measured in addition to an optogenetic mediated change in behavioral outcomes. This necessitates looking at conjunctions of success - cases in which there are both behavioral *and* histological effects or cases in which there are both behavioral *and* neurophysiological effects.

A second issue is that the unit of analysis for descriptive statistics in the original report is the single injection. It is perhaps not surprising that this database is organized injection-by-injection, given the success of single injections in rodent experiments to generate functional outcomes (Deisseroth, 2012). But, it is unclear whether single injections would have measurable outcomes in primates, particularly meaningful behavioral outcomes that are clearly grounded in neurophysiology and/or histology. Given that multiple injections of a viral vector into brain region are often required to achieve adequate coverage in larger NHP brains, a more appropriate basis to judge the relative success of experiments would include an analysis at the level of brain region and monkey and taking into consideration multiple markers of success simultaneously.

The summary of the resources in the paper by Tremblay states that optogenetics in NHPs has a 91% success rate (Figure 4D of Tremblay et al. 2020) - based on outcomes related to modulation in neural activity via histological outcomes measured in histology, *or* neurophysiology, *or* behavior, including injections where effects were weak or mixed. When only “strong” outcomes were considered, only 76% of injections were considered successful. In addition to being predicated on a broad definition of success, those rates are based on individual injections that may or may not have behavioral or physiological effects in NHP. Most importantly, the data points are clustered within animals, and it is animals, not neurons, which produce behaviors (Tinbergen, 1963).

Given the issues detailed above, we carried out a re-analysis of the database from Tremblay et al. (2020) with a specific focus on definitions of success that, in our opinion, are relevant to translational neuroscience for neuropsychiatric diseases – namely having both aneural AND behavioral outcome. We also used the experiment rather than the injection as the level of analysis. Further, we consider *where* the injection sites were in the brain and whether success (using our definition or the more liberal definition from Tremblay et al 2020) speaks to the readiness to use optogenetics productively in nonhuman primates to research or treat neuropsychiatric disease.

## Methods

We examined the publicly-available database described in the *Neuron* paper and then requested the unredacted spreadsheet from the first author so that we could unambiguously determine which experiments were performed in a single animal, something that was not possible using only the publicly posted database (https://osf.io/mknfu/). This allowed us to evaluate the number of successful experiments at the level of brain regions and animals, rather than single injections of viral vectors. In this way, we considered a single “experiment” to be one or more injections of the same viral vector into one brain area in one animal. Injections into two or more brain regions in the same monkey would be considered as two separate experiments in this analysis, even though they may be biologically constrained by being in the same animal. The choice of level of analysis in clustered data has a significant impact on outcomes, given that adjacent injections into the same area in the same animal could reasonably be considered to not be independent experiments (see also (Aarts et al., 2014). It is also this level of analysis that is most pertinent to behavioral neuroscience investigations of how neural circuits control behavior.

The database we received included 1085 injections. This was increased from the 1042 injections reported in the original publication, reflecting the utility of the database as an open resource (although it has not been updated since that time). We read this database into R and assigned experiment identifiers based on unique conjunctions of laboratory, animal identifier, and brain region, resulting in 383 unique experiments. We then read this database back out and carried out analyses to determine success rates at the experiment level. We categorized each injection as missing, failed, weak success, or strong success for histology, physiology, and behavior, attempting to follow the descriptions in Tremblay et al. 2020. Any experiment that had at least one injection classified as “strong success” was categorized as a strong success for that domain; experiments that had no strong success but at least one “weak success” were categorized as weak success, and so forth. Thus, this categorization may still overestimate the true success rate in cases where many injections were placed in a brain area but only a subset had successful effects. The redacted database with experiment identifiers and associated R analysis script are available at https://osf.io/j7rm2/.

## Results and Discussion

When we analyzed success rates considering injections as nested within monkey and brain region – what we refer to as individual “experiments” – rather than injection as the unit of analysis a somewhat different picture from that painted by Tremblay et al. (2020) emerged. First, very few experiments assessed all three outcomes (histology, *and* neurophysiology, *and* behavior). Only 39 out of the 383 cases we identified in the database (10.2%) evaluated the efficacy of an optogenetic manipulation in terms of histological, physiological, and behavioral effects. In this small subset of experiments there was a 59% (23/39) success rate on all three outcomes. The majority of these successes were in premotor cortex (7/9) and visual cortex (7/12). Put another way, only 6% (23/383) of the experiments attempted and reported were successful in using optogenetics to bridge levels of analysis from neurons to behavior.

When we examined each outcome domain separately, we found high success rates, as reported in the original paper, for histological (226/261, 86.6%) and neurophysiological (168/197, 85.3%) verifications of optogenetics. Success rates using behavioral outcomes (47/74, 63.5%) to validate optogenetic manipulations were lower. These success rates are lower than those reported and highlighted in the original paper, especially when we only considered strong effects: histology (72.4%), neurophysiology (75.1%), and behavior (52.7%). Moreover, these proportions consider only cases in which histology, physiology, or behavior was tested (Figure 2). The different success rates between histological and functional evaluations of optogenetics suggests the impediment in using optogenetics to study the primate brain is not merely viral transduction per se, but rather difficulties in modulating sufficient numbers of neurons in the larger primate brain (Gradinaru, 2020). Without question the further development of NHP specific genetic tools, rather than simply porting tools developed in rodent models to NHP, is necessary to meet the lofty goal of using NHPs as a translational bridge between behavioral and systems neuroscience and the clinic.

**Figure 2.**
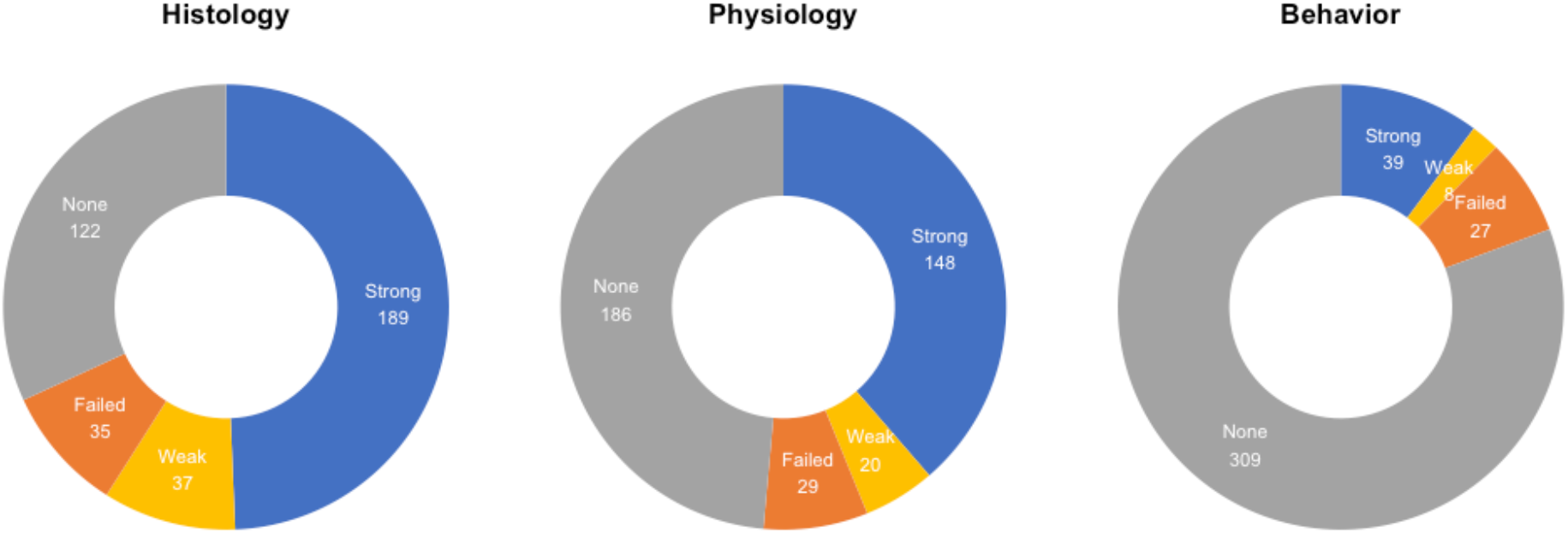
Numbers of experiments with histological, physiological, or behavioral outcomes indicating experiments that did not have/report a particular outcome (grey), cases where the experiment failed to detect effects (orange), cases where weak effects were detected (yellow), and cases where strong effects were detected (blue).

Speaking practically, a successful optogenetic experiment involves evaluating success in one or more of the outcome domains mentioned above. With the end goal of using optogenetics in NHPs to study the neural control of behavior at the level of neurons and circuits, it is reasonable to expect that behavioral manipulations should be validated in combination with either histology or neurophysiology. Therefore, we considered a definition of success as yielding a combination of effects (either behavior ⋂ neurophysiology or behavior ⋂ histology). Using a broad definition that included both weak and strong effects of success, there was a 62.5% (45/72) success rate among experiments that reported optogenetics effects on behavior and neurophysiology and a 62.5% (25/40) success rate among experiments that reported optogenetic effects on both behavior and histology. When we again adopted Tremblay and colleagues’ approach in considering only strong effects, success rates dropped to 48.6% (35/72) for studies that evaluated behavior in combination with neurophysiology and to 57.5% (23/40) for studies that evaluated behavior in combination with histology. Notably, these results are for experiments pooled across brain regions and laboratories. Perhaps more concerning for scholars interested in psychiatric and neurological diseases, is that 66.3% (254/383) of the experiments evaluated optogenetics in primary sensory or motor areas. Modulation of those areas should produce fairly stereotyped, clear functional (i.e., behavioral) outcomes, yet only 29 (11.4%; 29/254) of those experiments even evaluated both behavior and physiology. Cortex – particularly primate cortex – is heterogenous in its structure and success in one region does not guarantee success across the entire brain. Further, the brain’s inherent degeneracy and organization as a complex system means that different structures can accomplish, or contribute to, the same functional outcomes (Edelman & Gally, 2001) – as functional outcomes become more complex, so too do the possible configurations of neural circuity that can support them. This problem is compounded by the neocortical expansion of the primate brain (Wise, 2008). Practically, this means that the sorts of complex behaviors of interest to many NHP systems neuroscientists working are less likely be perturbed by the manipulation of single circuits.

In our view, a 62.5% (all effects) or 53.1% (strong effects only) success rate (averaged across the two conjunctions that include behavior as a functional outcome) is meaningfully different from the 91% (all effects) and 76% (strong effects only) success rate suggested by Tremblay et al. (2020). The lower success rate that we identified suggests that optogenetics is still in development as a tool in NHPs, rather than ready to be deployed to study functional outcomes in primates or to intervene in clinical contexts. This may reflect its true efficacy in mammals, previously masked by the lower costs and higher throughput associated with rodent models. Or, it may reflect issues with deploying viral techniques in primates – a perspective supported by mixed results of experiments using DREADDs (Designer Receptors Exclusively Activated By Designer Drugs) to manipulate behavior in primates, despite histological validation (Eldridge et al., 2016; Upright et al., 2018). Recognizing that further research is needed to determine the origin of the low success rates, it is nevertheless the case that at the moment success in obtaining strong effects across multiple functional outcomes when pursuing optogenetics in NHPs amounts to flipping a coin. This warrants a very different conclusion from the one that a reader might reasonably draw after reading the survey by Tremblay et al. (2020).

Even if we were to imagine that optogenetics was ready to be deployed in the way described by Tremblay and colleagues, there are a number of other major challenges that exist when translating the technology from use in rodents to use in primates, that must be considered. For example, for optogenetics to work, a light probe must be proximate to the structure being manipulated and this represents a significant engineering challenge in deep areas of the primate brain. That means that studies that demonstrate success in primary/secondary sensory or motor cortex which sit at the surface of the brain are fundamentally different from those evaluating the manipulation of structures deep within the brain. Accessing deep structures – for example the amygdala, hippocampus, and subgenual cingulate cortex – that are part of circuitry relevant to psychiatric and neurological disorders without damaging the areas through which probes are passed is a significant engineering challenge that has not yet been sufficiently addressed. Moreover, when optogenetics was successfully used to perturb a deep structure, like the superior colliculus (Cavanaugh et al., 2021), it took advantage of the detailed functional mapping that specified how the superior colliculus controls saccadic eye movements. We are nowhere near such refinements in knowledge when we consider neural circuits relevant for treating psychiatric disease. We are a long way from conquering the engineering problems, much less interpretive issues, that will allow for the “development of clinical technologies relying on optogenetics to control neural populations and pathways with unprecedented precision” as the authors claim (p. 12-13). It is shortsighted to assume that translation will work when so many basic experiments targeting areas with known anatomical structure and functions fail. When optogenetics sufficiently matures to be used as a reliable tool in primate systems neuroscience, it is more likely to allow elegant circuit dissections of behavior that can inform existing treatments than to be a frontline treatment for psychiatric disorders (Lüscher & Pollak, 2016).

### Where is nonhuman primate optogenetics in the hype cycle?

As the drive for new technology pushes forward, we propose that the ability to manipulate behavior using optogenetics (or other genetic tools, such as DREADDs) should remain the gold standard by which we evaluate its success. This view echoes recent calls to reexamine the utility of behavior itself in neuroscience (Krakauer et al., 2017; Niv, 2020). We would suggest that an approach with a current success rate of 53.1% or 62.5% (on average) should be evaluated differently than one with a success rate of 76% or 91% in terms of choosing a method for interrogating function of neural circuits. Given this, we propose that more effort needs to be devoted to validating and optimizing optogenetics and related technologies specifically in NHPs. Validation and optimization is needed before such tools can be efficiently deployed in experimental contexts, and certainly before claims about their translational relevance are made. Such basic science efforts may not have the glamour or appeal of new technological invention or the deployment of new technology in experimental contexts, but they are vital for moving the field forward and ultimately for generating new effective treatments and interventions of psychiatric and neurological disease.

Given the data in the open resource, we propose that the state of the technology for use in NHPs may be oversold – that is, it is being discussed in the literature as if it is in the plateau of productivity, but in reality it is much earlier along the hype cycle. This has significant consequences for what work is funded and can be carried out and creates a tension between hypothesis-driven work that speeds scientific discovery and the need to employ methods that are unreliable in NHPs. The drive for technological innovation from the BRAIN Initiative (https://braininitiative.nih.gov/) has created a primacy of technology such that optogenetics and related methods are often viewed by science funders as a better approach for systems neuroscience than “classical” experimental methods in spite of interpretive difficulties with transient manipulations of neural activity (e.g., Otchy et al., 2015).

Believing that a technology is further along in the hype cycle than it actually is may result in abandoning tools that actually work before the new tools are actually ready. This can limit what science is done and the inferences that can be drawn from that science. Imagine, for example, using such a new technology to temporarily deactivate a hub of the primate brain thought to be involved in neuropsychiatric disease, about which there are functional hypotheses but for which the function has not been clearly established. A scientist deploys the technology, measures behavior, and gets null results. Without established ground truth about the brain area’s function (determined with the use of “old” tools), there is no way to determine if the functional hypothesis was wrong or whether temporary deactivation leads to different behavioral consequences that were observed in permanent deactivation, stimulation, or recording studies (the “old” tools), or if the technology simply didn’t work. Until the function of the tool is well established, it is critical that we maintain tried and true methods (Katz et al., 2016; Vaidya et al., 2019), deployed in the context of testing interesting ideas to both speed scientific discovery about the brain but also to help us improve the functionality of the new tools.

### What is the way forward?

The data provided by the open NHP optogenetics resource leads us to conclude that optogenetic manipulation is not yet at a state of development where it can be reliably applied to investigate functional effects of neural circuit manipulations in NHPs. Major questions and challenges remain about the application of this technology, both in terms of methodological approaches (e.g., how to target deep structures without causing significant brain damage, how to achieve transfection coverage of those structures at a rate sufficient to impact behavior, etc.) and theoretical ones (e.g., what is positive evidence that the technology is ready to move into humans, let alone be deployed in greater numbers of NHPs). In contrast to being a mature approach that is ready for translation in humans, we suggest that as a technology it is much earlier along the hype cycle than one might believe at first blush – perhaps even at inflection following the “innovation trigger”. We hope that critical considerations of where we are as a field, like this one, might move us as a community to the “peak of inflated expectations” allowing us to move the technology forward along the cycle. We celebrate the meta- and open science nature of the database, as well as the efforts of project leader and the individual scientific teams, because they allow for a gestalt evaluation of the efficacy of optogenetics in the NHP brain, which we hope will ultimately speed science and discovery.

We suggest, based on our re-analysis of the data in the open NHP optogenetics resource, that this technology is still immature, with few successes in modulating circuits that are relevant for psychiatric neuroscience. We do <u>***not***</u> suggest, on this basis, that the enterprise of developing optogenetic approaches in NHPs should be abandoned. But, we would propose that perhaps more fundamental engineering and validation work should be a priority to increase the success rate of this approach, especially in brain structures and circuits outside of primary sensory and motor function. We would also suggest that as systems and cognitive neuroscientists, we reflect on the relationship between technological innovation and fundamental questions that are critical to our field, as well as their translation to psychiatry. The ability to execute technologically dazzling “proof of principle” experiments potentially comes at a cost of allocating resources to research that uses less technologically advanced approaches but nevertheless is effective in addressing fundamentally critical questions. As we have argued here and elsewhere (Baxter and Costa, 2021), the interpretation of the functional impact of any manipulation of neuronal activity cannot come from a single technical strategy. Especially for translational research in NHPs, innovation in relating circuits to behavior with the potential for elucidating mechanisms of psychiatric disease cannot be hobbled by reliance on as-yet-unstable technological approaches.

## Acknowledgements

The authors wish to thank Peter Rudebeck for helpful comments on a previous draft of this manuscript.

